# BDQC: a general-purpose analytics tool for domain-blind validation of Big Data

**DOI:** 10.1101/258822

**Authors:** Eric W. Deutsch, Roger Kramer, Joseph Ames, Andrew Bauman, David S. Campbell, Kyle Chard, Kristi Clark, Mike D’Arcy, Ivo D. Dinov, Rory Donovan, Ian Foster, Benjamin D. Heavner, Leroy E. Hood, Carl Kesselman, Ravi Madduri, Huaiyu Mi, Anushya Muruganujan, Judy Pa, Nathan D. Price, Max Robinson, Farshid Sepehrband, Arthur W. Toga, John Van Horn, Lu Zhao, Gustavo Glusman

**Author notes:** Correspondence, Gustavo Glusman, Institute for Systems Biology, 401 Terry Ave N, Seattle, WA 98109, USA. equal contribution (Eric W. Deutsch, Roger Kramer).

## Abstract

Translational biomedical research is generating exponentially more data: thousands of whole-genome sequences (WGS) are now available; brain data are doubling every two years. Analyses of Big Data, including imaging, genomic, phenotypic, and clinical data, present qualitatively new challenges as well as opportunities. Among the challenges is a proliferation in ways analyses can fail, due largely to the increasing length and complexity of processing pipelines. Anomalies in input data, runtime resource exhaustion or node failure in a distributed computation can all cause pipeline hiccups that are not necessarily obvious in the output. Flaws that can taint results may persist undetected in complex pipelines, a danger amplified by the fact that research is often concurrent with the development of the software on which it depends. On the positive side, the huge sample sizes increase statistical power, which in turn can shed new insight and motivate innovative analytic approaches. We have developed a framework for Big Data Quality Control (BDQC) including an extensible set of heuristic and statistical analyses that identify deviations in data without regard to its meaning (domain-blind analyses). BDQC takes advantage of large sample sizes to classify the samples, estimate distributions and identify outliers. Such outliers may be symptoms of technology failure (e.g., truncated output of one step of a pipeline for a single genome) or may reveal unsuspected “ signal” in the data (e.g., evidence of aneuploidy in a genome). We have applied the framework to validate real-world WGS analysis pipelines. BDQC successfully identified data outliers representing various failure classes, including genome analyses missing a whole chromosome or part thereof, hidden among thousands of intermediary output files. These failures could then be resolved by reanalyzing the affected samples. BDQC both identified hidden flaws as well as yielded new insights into the data. BDQC is designed to complement quality software development practices. There are multiple benefits from the application of BDQC at all pipeline stages. By verifying input correctness, it can help avoid expensive computations on flawed data. Analysis of intermediary and final results facilitates recovery from aberrant termination of processes. All these computationally inexpensive verifications reduce cryptic analytical artifacts that could seriously preclude clinical-grade genome interpretation. BDQC is available at https://github.com/ini-bdds/bdqc.

## Background

As data volumes increase and analysis pipelines become more complex and more expensive to (re)run, validating data, both the input to a pipeline and its output, becomes more critical. Recently, several tools have become available to perform quality control (QC) of ‘omics data, with an emphasis on high-throughput sequencing data. BatchQC [1] is a toolkit that focuses on identification and correction of batch effects in genomics data. AlmostSignificant [2] simplifies the aggregation of many QC metrics with metadata, for the evaluation of the voluminous data from DNA sequencing runs. MultiQC [3] flexibly integrates multiple QC tools into a single visual report spanning multiple samples and revealing trends and biases. These and similar tools commonly focus on the QC of the original data sets—e.g., the output of sequencing machines—that serves as initial input to analysis pipelines. Of necessity, these QC tools tend to be very domain-specific, with algorithms and visualizations that are tailored to the data types typically observed in genomics and multi-omics research.

Considering the complexity of modern analysis pipelines, we recognize the additional need for QC at later stages in analysis. Data analysis pipelines typically undergo development concurrently with the research that depends on them. Validating these pipelines is essential to ensure the integrity of research results. While software validation is a standard component of the development process, both “white box” (e.g., unit testing) and “black box” testing (e.g., of a complete software package) are typically limited to confirming the expected transformations of exemplary inputs. Validating data usually means confirming that expected regularities exist or, conversely, identifying departures from expected regularity.

Considering the central characteristics of Big Data—large cardinality and complexity of data elements—we also recognize the opportunity for applying data analysis techniques to QC purposes. Furthermore, such application need not be informed by domain-specific knowledge. There is, on deeper consideration, not a significant distinction between data validation and data analysis. Data analysis typically involves relating two or more bodies of data, but identifying regularity and departures from it are inherent in both validation and analysis. The primary difference between validation and general data analysis resides in the resolution of expected regularities. Concretely, distributions are typically narrower in the context of validation. However, a spectrum of analyses are possible on any given body of data from merely characterizing its representational attributes to progressively deeper examination of its content. Statistical expectations will exist at every level, and it is not likely useful to try to establish a clear boundary between “mere” validation and general analysis.

*Example*. If a pipeline is executed on multiple similar inputs, all output files may be expected to be of approximately the same size or contain approximately the same number of lines (assuming they are line-oriented text). If the pipeline is, for example, a whole-genome analysis and input is *Homo sapiens* sequence or variant-call data, a “chromosome”column may be expected to contain exactly 24 or 25 discrete labels, depending on presence and treatment of the allosomes and mitochondria (i.e., 1̤22, M, X and potentially Y). Fewer labels—the software need not even “know”these are chromosome labels—may indicate premature pipeline termination—that is, a runtime error. It is notable that validation need not be concerned with the exact content of the chromosome labels—for example, whether rows with mitochondrial features are labeled “MT” or “chrM.” Furthermore, one need not know the expected count a *priori*. It may be sufficient to confirm that the corresponding columns in all of a set of files contain the same counts of discrete tokens. These considerations suggest that it should be possible to create a general-purpose data validation software tool, “general purpose” in the sense that its design does not need to be tied to any specific knowledge domain or even data format. To the extent that it confines its function to the representation of data itself, not the phenomena described by data, it should be applicable to any data. This need not preclude data and domain specific features; any implementation should explicitly support “extensibility” to push as deeply into content-aware validation as is sensible with respect to usability considerations.

We present here a framework for Big Data Quality Control (BDQC): an extensible set of analyses, heuristic and statistical, that identify deviations in data without regard to its meaning (domain-blind analyses). BDQC takes advantage of large sample sizes to analyze the files, estimate distributions and identify outliers. Such outliers may be symptoms of technology failure (e.g., truncated output of one step of a pipeline for a single genome) or may reveal unsuspected “signal” in the data (e.g., evidence of aneuploidy in a genome).

## Results

### A general-purpose method for detecting file anomalies

We have created a framework and tool for Big Data Quality Control (BDQC). Given a collection of files, which could be input data, intermediate results in a pipeline, analysis results, log files, etc., BDQC evaluates the structure and content of the files to create data-driven models, then evaluates each file vs. the models to identify outliers. Files may be outliers for a variety of reasons, including misclassification, technical failures during production or transmission, batch effects, etc.

BDQC analyzes a collection of files in three stages (Figure 1); these files are assumed to be of the same type. First, BDQC analyzes each file individually and produces a summary of the file's content (Within-file Analysis). Second, the aggregated file summaries are analyzed heuristically (Between-file Analysis) to model the expected content of the files. Finally, individual file summaries are compared with the models (Evaluation) to identify possible anomalies. A file is considered “anomalous” when one or more of the statistics computed on its content (Within-file Analysis) are outliers relative to the corresponding model. The three stages of operation can be run independently. In particular, models derived from one set of files can be reused to evaluate other files that were not used for training the model, as long as they are of the same type as the files in the initial training set.

**Figure 1.**
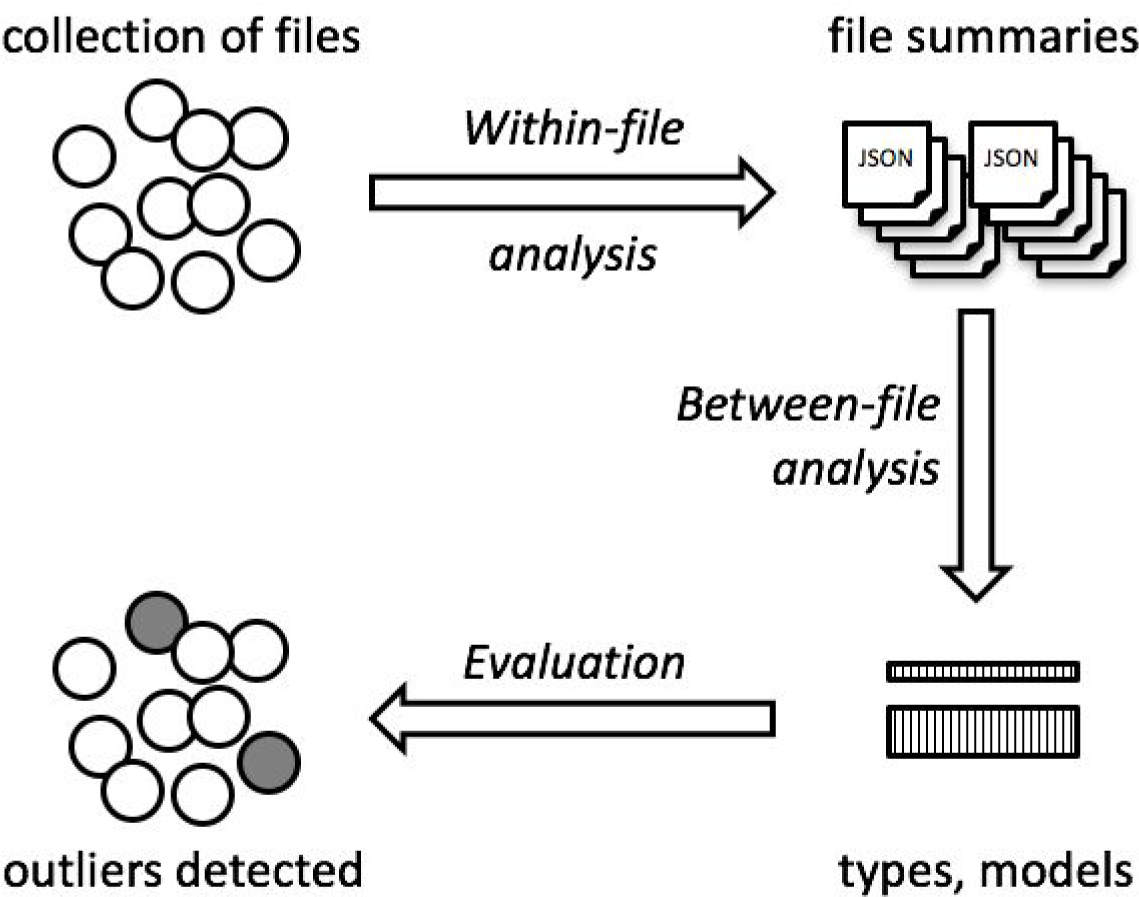
Overview of the method. A collection of files, ideally of the same type, are studied individually (Within-file analysis) yielding file summaries in JSON-format. Multiple such file summaries are combined and modeled (Between-file analysis) yielding a series of content models. Finally, these models can be used to evaluate files to detect anomalies (outliers).

In the Within-File analysis step, the BDQC method uses a framework that is extensible via plugins (Figure 2). Out-of-the-box, the framework includes a small number of general-purpose plugins for identifying file properties like filename, file size and modification time, and for the more detailed analysis of files with tabular format. Domain-specific plugins can be developed for achieving higher sensitivity of outlier detection for specialized file formats. Each plugin may compute multiple statistics for each file. The Between-File analysis step collates these statistics from multiple files into a summary matrix, which is then studied to derive models and identify outliers.

**Figure 2.**
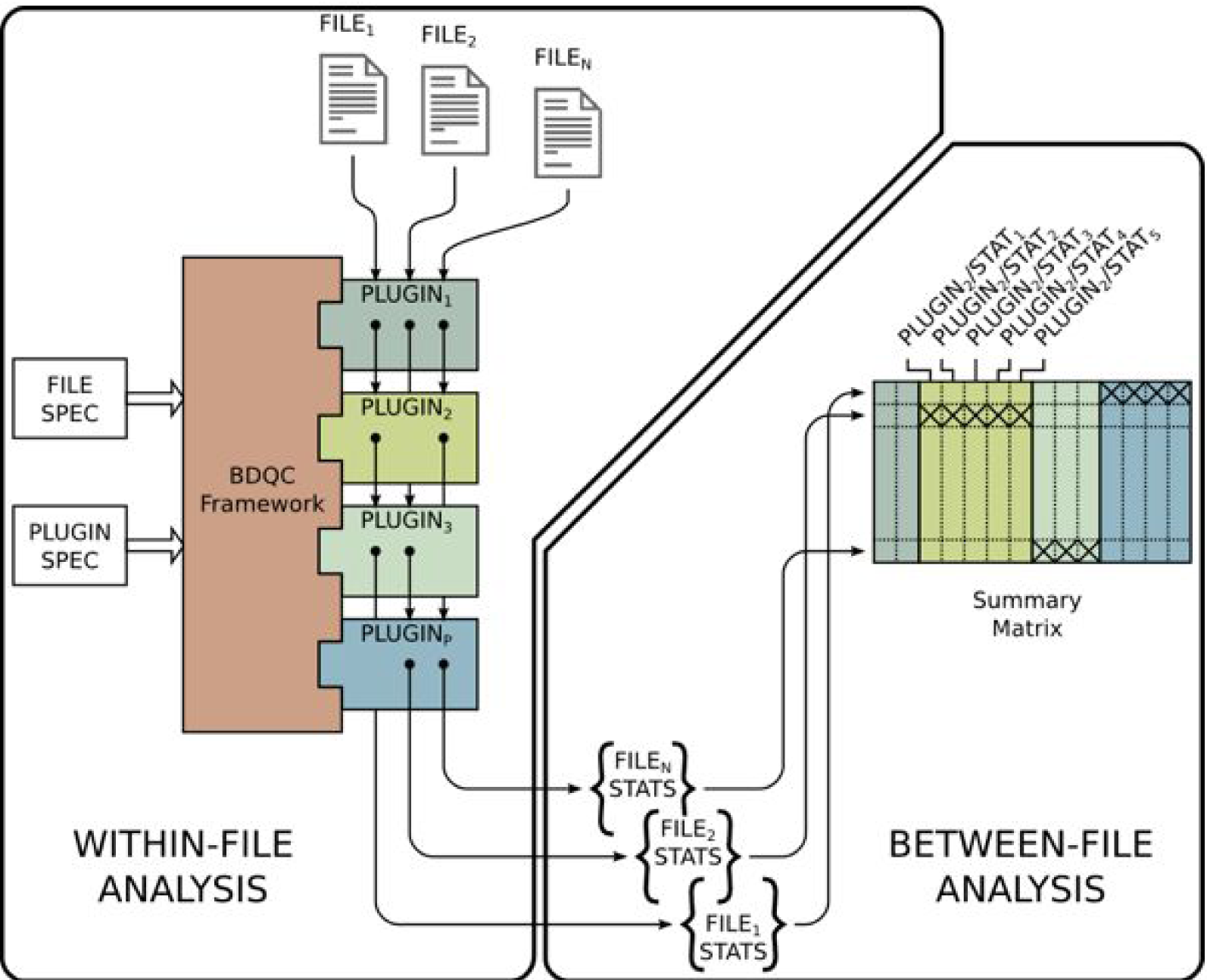
More detailed representation of the BDQC framework, in which plugins compute multiple statistics that are collated into a matrix for modeling. Not all plugins need to be executed on all files: plugin #3, for example, was not run on file #N.

### QC of a small data set

We obtained analytical results of differential expression from RNAseq data of the Accelerating Medicines Partnership-Alzheimer’s Disease (AMP-AD) project and studied 26 files from the MayoRNAseq study [4]. For each of two tissues (cerebellum and temporal cortex), the data set includes 12 pairwise comparisons and one gene annotation/conditional quantile normalization file (Table 1). We performed seven tests including different subsets of the 26 files. In a first exploratory test, we studied all 26 files together. This test clearly identified the two ‘GeneAnnotation_cqnCounts’ files as outliers, with over 40 flags each. These two files indeed have different structure and content from the rest of the data set - a fact easily comprehensible to a human but automatically detected here without prior knowledge of the data set. Testing cerebellum and temporal cortex files separately likewise identified the ‘GeneAnnotation_cqnCounts’ files as outliers. Excluding these outlier files from the test, and other combinations of pairwise comparison files (Simple and Comprehensive) raised few flags, suggesting that the data set is well curated. Nevertheless, all tests including the file labeled ‘CBE_CONvsPA_DEG_Simple’ raised a few flags, suggesting potential quality issues with this file. More detailed examination of the BDQC output revealed that the ‘Dx.qValue’ column of this file takes only three possible numerical values, while other files in the data set have many thousand possible values in the corresponding column.

**Table 1.** Outlier detection in the AMP-AD MayoRNAseq differential gene expression data set for cerebellum (CBE) and temporal cortex (TCX). Columns represent tests including different subsets of the files: All: all files, CBE: cerebellum only, TCX: temporal cortex only, CBE-: cerebellum pairwise comparisons, TCX-: temporal cortex pairwise comparisons, Simple: all files labeled ‘Simple’, Comp: all files labeled ‘Comprehensive’. OK: included in the test and raised no flags. Empty cells in the table denote files not included in each test. Files labeled ‘GeneAnnotation_cqnCounts’ are recognized as outliers with multiple flags (orange background) relative to the pairwise comparisons between Alzheimer’s Disease (AD), progressive supranuclear palsy (PSP), pathologic aging (PA) and elderly controls (CON).

| File name | File size | All | CBE | TCX | CBE- | TCX- | Simple | Comp |
| --- | --- | --- | --- | --- | --- | --- | --- | --- |
| CBE_CONvsAD_DEG_Comprehensive.txt | 9178381 | OK | OK |  | OK |  |  | OK |
| CBE_CONvsAD_DEG_Simple.txt | 9200164 | OK | OK |  | OK |  | OK |  |
| CBE_CONvsPA_DEG_Comprehensive.txt | 10618075 | OK | OK |  | OK |  |  | 1 |
| CBE_CONvsPA_DEG_Simple.txt | 10592279 | 1 | 2 |  | 2 |  | 3 |  |
| CBE_CONvsPSP_DEG_Comprehensive.txt | 9209392 | OK | OK |  | OK |  |  | OK |
| CBE_CONvsPSP_DEG_Simple.txt | 9222336 | OK | OK |  | OK |  | OK |  |
| CBE_PAvsAD_DEG_Comprehensive.txt | 9603109 | OK | OK |  | OK |  |  | 1 |
| CBE_PAvsAD_DEG_Simple.txt | 9608155 | OK | OK |  | OK |  | 1 |  |
| CBE_PAvsPSP_DEG_Comprehensive.txt | 9689684 | OK | 1 |  | OK |  |  | OK |
| CBE_PAvsPSP_DEG_Simple.txt | 9722360 | OK | OK |  | OK |  | OK |  |
| CBE_PSPvsAD_DEG_Comprehensive.txt | 8970854 | OK | OK |  | OK |  |  | OK |
| CBE_PSPvsAD_DEG_Simple.txt | 8961051 | OK | OK |  | OK |  | OK |  |
| CBE_GeneAnnotation_cqnCounts.txt | 33628279 | 41 | 51 |  |  |  |  |  |
| TCX_ADvsCON_Comprehensive.txt | 9211843 | OK |  | OK |  | OK |  | OK |
| TCX_CONvsAD_DEG_Simple.txt | 9290169 | OK |  | OK |  | OK | OK |  |
| TCX_CONvsPA_DEG_Comprehensive.txt | 10763514 | OK |  | OK |  | OK |  | 1 |
| TCX_CONvsPA_DEG_Simple.txt | 10760142 | OK |  | OK |  | OK | 1 |  |
| TCX_CONvsPSP_DEG_Comprehensive.txt | 9321440 | OK |  | OK |  | OK |  | OK |
| TCX_CONvsPSP_DEG_Simple.txt | 9326364 | OK |  | OK |  | OK | OK |  |
| TCX_PAvsAD_DEG_Comprehensive.txt | 9674145 | OK |  | OK |  | OK |  | OK |
| TCX_PAvsAD_DEG_Simple.txt | 9705354 | OK |  | OK |  | OK | OK |  |
| TCX_PAvsPSP_DEG_Comprehensive.txt | 9775332 | OK |  | OK |  | 2 |  | 1 |
| TCX_PAvsPSP_DEG_Simple.txt | 9805463 | OK |  | OK |  | 1 | 1 |  |
| TCX_PSPvsAD_DEG_Comprehensive.txt | 8950644 | OK |  | OK |  | OK |  | OK |
| TCX_PSPvsAD_DEG_Simple.txt | 9034496 | OK |  | OK |  | OK | OK |  |
| TCX_GeneAnnotation_cqnCounts.txt | 24734242 | 43 |  | 60 |  |  |  |  |

### QC of an analysis pipeline

We studied intermediate files in our previously published coverage analysis pipeline for WGS data [5]. Briefly, the pipeline condenses depth-of-coverage information into a compact format, computes summary statistics, computes a Reference Coverage Profile (RCP) from multiple such files, normalizes each genome’s coverage relative to the RCP, uses a hidden Markov model (HMM) to segment the normalized coverage into regions of uniform ploidy, and filters these results to identify regions of unusual ploidy relative to a reference population (Figure 3). Here, we used BDQC to evaluate the output files of four steps in this pipeline (green arrows in Figure 3). We studied a large collection of 4461 genome assemblies produced by Complete Genomics, Inc. using a wide variety of software versions as described [5], flavors of the technology, and two reference versions (hg18 and hg19). We visualized the results (Figure 3) and observed two types of failure: missing files and BDQC-flagged outliers. In some cases the absence of output files was expected, since we had not executed the pipeline for genome assemblies on the hg18 reference. In other cases, we observed pipeline failure: early stages of the pipeline were completed but later stages were not. Interestingly, BDQC raised multiple flags (denoting outlier values, red marks in Figure 3) for the early stage output files that preceded missing late stages. This indicates that while those early stage analysis steps produced an output, the results were deficient or compromised in ways that caused subsequent pipeline steps to fail. In several of these cases we could verify the presence of specific analytic failures, e.g., premature termination leading to truncated output files, which could be rectified by repeating the analysis on those specific samples. Such interventions led to successful pipeline completion.

**Figure 3.**
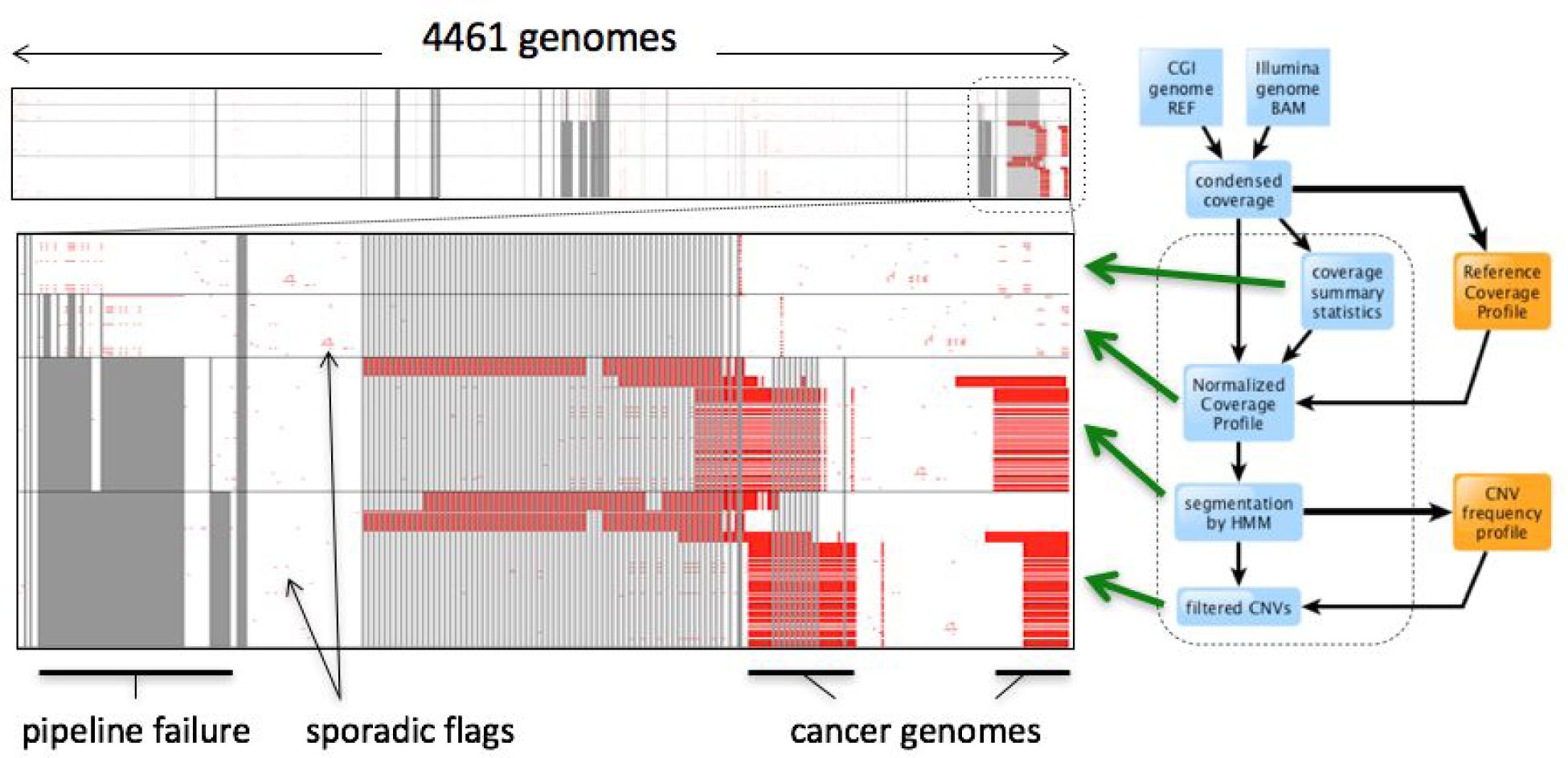
BDQC applied to the intermediary files in a genome coverage analysis pipeline, depicted on the right side. Upper left: overview of results over 4461 genomes; lower-left: expanded view of a subset of genomes. Red dots represent outlier flags; gray lines represent missing files.

We also observed sporadic flags, most of which likely represent content deviations in the data and/or batch effects. Finally, we observed multiple genomes with some flags for early pipeline stages (summary statistics and normalized coverage) and large numbers of flags for later pipeline stages (segmentation and filtering). Most of these genomes are derived from cancer samples; these indeed have unusual ploidy patterns relative to germline-derived genomes,which represent the majority of the samples studied here.

## Discussion

We presented here BDQC, a framework and tool for automated, domain-agnostic quality control of Big Data. The framework evaluates the structure and content of the files, progressing from identification of file type and structure to modeling of their content. Based on the foundational concept that “sanity checks” can usually be carried out using summary statistics that are blind to the meaning of the content, BDQC creates data-driven models, then evaluates each file vs. the models to identify outliers. Files may be outliers for a variety of reasons, including misclassification, technical failures during production, transformation or transmission through a network, batch effects, and more.

We developed the BDQC framework with several explicit goals in mind:

1. Identify “anomalous” files among a large collection of similar files of arbitrary type with as little guidance from the user as possible - ideally none. In other words, it should be useful “out of the box” with almost no learning curve.
2. “Simple things should be simple; complex things should be possible”. Although basic use should involve almost no learning curve, it should be possible to extend it with arbitrarily complex (and possibly domain-specific) analysis capabilities. We achieve this by implementing an architecture that can be extended via plugins.
3. Plugins should be simple to develop, and the framework should be robust to faults in plugins. BDQC’s plugin system supports the development of Perl and Python plugins that can implement the analysis code directly, and can also be used to wrap existing stand-alone analysis tools programmed in any language.
4. The result of the QC should be informative about the location and nature of the failure. Since the metrics being modeled in BDQC typically refer to specific aspects of the file structure (line count, delimiters, the content of specific columns, etc.), the location of the failure is straightforward to identify, and the specific details of what makes the file an outlier for that metric are typically informative about the nature of the failure (e.g., unusual values observed).

For BDQC analysis to yield meaningful results, the input files need to be comparable, i.e. ‘of the same type’. As we demonstrated in the MayoRNAseq example, if a small number of files of a different type are present in a data set, they are readily identified as outliers; their exclusion increases the sensitivity of analysis of the remaining files. As currently implemented, BDQC will be less successful if confronted with a collection of files of disparate types in comparable proportions. Future versions of BDQC will include an internal type classification step that will enable automated analysis of complex, mixed data sets.

## Methods

### Implementation

The BDQC tool is implemented in the Perl programming language; we also provide an extensible framework for performing Within-file Analysis, implemented in Python3 and with critical sections implemented in C for efficiency. This framework is extensible via a modular plugin architecture. The BDQC protocol may be invoked via the command line interface. Command line arguments control which files are studied, how files are summarized, and how the summaries are aggregated and finally analyzed. All command line arguments are optional; the framework will carry out default actions.

A successful BDQC execution ends with one of 2 general results:

1. No outliers were found.
2. Anomalies (‘flags’) were detected in specific files. In this case, a report is generated summarizing the evidence, as text or optionally as an interactive visualization.

### Within-file Analysis

The plugins that are executed on a file entirely determine the content of the summary (the statistics) generated for that file. The BDQC framework:

1. assembles a list of paths identifying files to be analyzed,
2. executes a *dynamically-determined* subset of the available plugins on each file path,
3. merges the plugins' results into one (JSON-format) summary per analyzed file.

In the extensible Python framework, each plugin can declare (as part of its implementation) that it depends on zero or more additional plugins. The extensible Python framework performs the following actions:

1. ensures that a plugin's dependencies execute before the plugin itself
2. provides each plugin with the execution results of its *declared* dependencies
3. minimizes work by only executing a plugin when required (Figure 2)
4. optionally stores the summary for file foo.txt in an adjacent file named foo.txt.bdqc

### Plugins

The BDQC executable *framework* does not itself examine files' content. All *Within-file* analysis is performed by plugins. Several plugins are included in (but are, nonetheless, distinct from) the framework. These plugins, referred to as “Built-ins”, provide very general purpose analyses and assume *nothing* about the files they analyze. Although their output is demonstrably useful on its own, the built-in plugins may be viewed as a means to “bootstrap” more specific (more domain-aware) analyses.

Plugins provide functions that can read a file and produce one or more summary statistics about it. The functions are expected to implement a standard interface, and the plugin is expected to export certain symbols used by the BDQC framework.

A Python plugin has several required and optional elements shown in the example below.

~~~
VERSION=0×00010000
DEPENDENCIES = [‘bdqc.builtin.extrinsic’,‘some.other.plugin’]
def process( filename, dependencies_results ):
    # Optionally, verify or use contents of dependencies_results.
    with open( filename ) as fp:
        pass # …do whatever is required to compute the values
    # returned below…
    return {
        ‘a_quantitative_statistic’:1.2345,
        ‘a_3x2_matrix_of_float_result’:[[3.0,1.2],[0.0,1.0],[1,2]],
        ‘a_set_result’:[‘foo’,‘bar’,‘baz’],
        ‘a_categorical_result’:“yes” }
~~~

### Between-file Analysis

After individual signature metrics are collated for all files of one class, the vectors of values are passed individually to the modeling and outlier detection component of BDQC. Each vector of values is analyzed separately to create a model of what is typical for that metric for that collection of files. Null values are first removed (although considered further, as described below), and then the vector of numerals is sorted and such statistics as mean, median, standard deviation, and quartiles are calculated. Consider an example where 100 plain text files make up a class, wherein each file was analyzed with the Text plugin. One of the many metrics that are produced is the metric for number of lines in the file, which varies among the 100 files analyzed. In this case a vector of 100 integers is passed to the modeling component to determine what is typical and which elements are outliers. The return of the modeling component is a model of typical that can be stored for later as well as a 100-element vector in which several values are stored for each input datum, including the original value, the degree of deviation from the typical value, and whether the datum is an outlier.

#### Modeling

A simple way to calculate a model would be to compute a mean and standard deviation for a distribution of numbers and then label any data points more than N standard deviations from the mean an outlier. This does not work well for distributions with outliers as the computed values can be skewed by the outliers. A calculated median and semi-interquartile range (SIQR) is less sensitive to outliers, but may also not work well with larger populations or ones with non-Gaussian tails. For BDQC we do compute these values and report them, but also implement an outlier detection algorithm that finds the largest gap between a sorted list of data points, both at the top end and the bottom end separately to account for skewed distributions. If the largest gap is at least twice the size of the next largest gap, and if that largest gap is inside the outer quartile, then all points beyond this are considered outliers. Then for each data point, a deviation value is calculated based on the gap size. This approach seems to replicate what the human mind will consider outliers.

While the above approach works well for many-valued distributions, it does not work well for two-valued distributions (i.e. a vector with only two possible values), so there is special code that arbitrarily determines how many of the smaller population are permitted before both values are considered normal. For example, in a set of 50 data points, if two were 14 and 48 were 15, the two 14s would be considered outliers. However, is there were 25 14s and 25 15s, then both would be considered normal. These rules are somewhat arbitrary, but seem to work well.

Vectors of string type are handled in a similar way, except that the strings are first converted to numerals. The conversion is simply the linear combination of the length of the string and the average ASCII value of the characters in the string. Thus, longer strings increase the numeral and higher ASCII values (such as lower case letters over upper case letters) increase the numeral. The distribution of numerals produced in this way is also modeled as a set of numbers as described above and outliers are detected as above. This easily separates most kinds of deviations from typical values.

#### Handling of nulls

An exception to this scheme is the handling of null (absent) values. When a vector contains nulls, those nulls are first removed and the model is created as described above based on the remaining non-null values. Outliers for non-null values are computed as described above. Then, the number of nulls in the distribution dictates whether the nulls are outliers. A small number of nulls will be considered outliers, while a small number of non-nulls will be considered the outliers, and within those bounds, null will be considered part of normal. For example, in our example above with a vector of 100 values: if there are only 2 nulls, then those nulls are outliers in addition to any outliers computed from the distribution of 98 integers. If 98 of those values are nulls and only two are integers, then null is considered normal and the two integers are the outliers. Finally, if there are 50 nulls and 50 integers, then outliers are computed from the distribution of integers and nulls are considered a part of normal. The boundaries between these regimes are set arbitrarily to something that seems reasonable, varying with the size of the population.

#### Handling of complex data types

The above description applies to the handling of vectors of any scalar data type. Two additional complex data types are supported, histograms and arrays. The histogram data type is set of discrete values of any type and an associated number (typically a count). Metrics that are expected to scale with the file size in the same proportions make for good histograms. For example, for a binary file the overall byte counts are put in a histogram, and for an XML file the overall counts for each element name are put in a histogram. One generally expects a file twice as large to have more counts of each element with a constant proportion. Internally to BDQC, all histograms in the vector for a given metric are averaged to create an average histogram normalized to unity (as the union of all values in any histogram, with missing values as zeros). Then, the variance of each individual histogram (normalized to unity) is computed against the average histogram. The resulting vector of variances is then modeled as above like any other scalar vector and outliers detected.

Arrays are somewhat similar to histograms in that there is a list of discrete values, but the difference is that there is no count for each value. Arrays are modeled differently in order to accentuate rare events. First the union of all values across all files is assembled, along with the count of how many files each value is found in. In order to accentuate values found in only 1 or a small number of files, each count is transformed with a[i] = exp(-1 * ((count[i]-1) / maxcount * 10 + 1)) and the scalar numeral for each file is the sum of a[i] for all observed values i. This ensures that a file that has a very rare value in it will get a higher value than files with no rare values. The resulting vector of scalar summations is then modeled as above like any other scalar vector and outliers detected. Consider an example of a column of data that contains thousands of instances of 25 possible discrete values, perhaps “chr1”, “chr2”, etc. If most of these files contain many instances of each, but one file also contains a few instances of “chrM”, then this new value in the list of one file would cause a radically different value for that file, which would be easily flagged by the outlier detection algorithm. Of course, if many files each had a value unique to it, then having a unique value would be considered typical and not be flagged.

#### Model export and reuse

Each of these models for each signature attribute is serialized to a models file. Future version of BDQC will allow signature calculation for an additional (presumably small) set of files and then direct comparison to these persisted models. Therefore it would easy and quick to classify and detect outliers among new files by comparing to models previously trained on thousands of files.

### Visualization of results

The generic endpoint output of invoking BDQC is a list of outlier files and flags that were raised to put them on the outlier list. This list is generally returned as a text report that is easily read by the user. For integration into pipelines, the data may be optionally returned as a tab-separated value table or as a JSON object. Finally, the output can also be written in HTML as an interactive web interface that depicts all the outliers within each file type and an interactive plotting window implemented using Plot.ly [6]. The plotting window enables users to view the numerical distribution of each vector for each signature attribute, with outliers flagged as red data points. Mouse-over functionality shows the filename associated with each data point. The usual zooming and panning functionality of the Plot.ly toolkit is enabled to allow for detailed human inspection of the results. See Figure 4 for example with the MayoRNAseq dataset.

**Figure 4.**
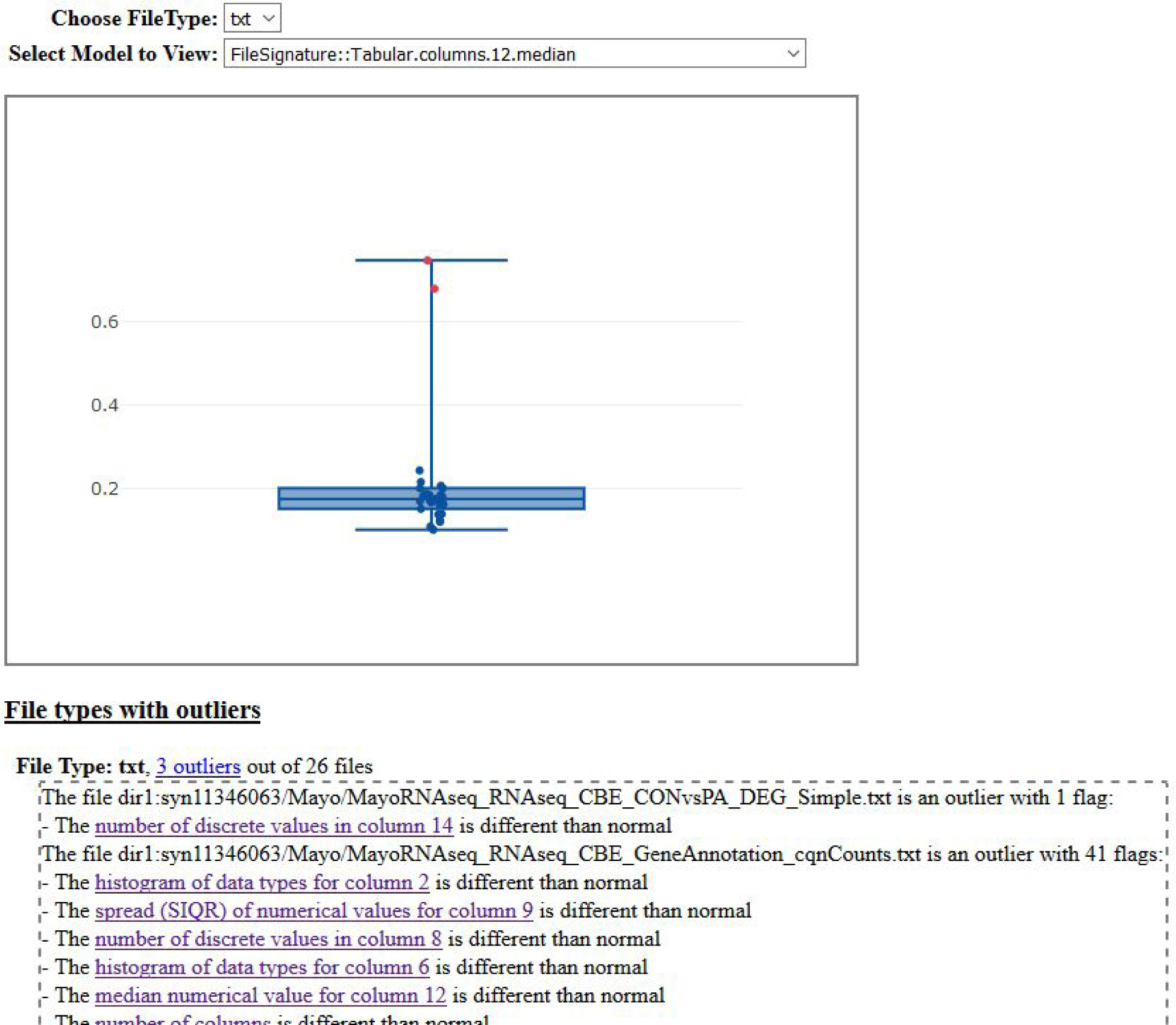
Example web browser-based visualization of a small data set. In this example, there is a single type of file, a “txt” type based on the file extension. The BDQC framework automatically determined that they are tab-separated value files, and analyzed them with the Tabular plugin. There are three outlier files identified among the 26 files. Clicking on a hyperlink associated with each flagged metric displays the distribution of data points collect from the files for that metric. In this example, clicking on the “median numerical value for column 12” hyperlink displays this metric for all files, with the outliers colored in red.

### Validation Data

#### RNAseq differential expression data

We obtained analytical results of differential expression from RNAseq data of the Accelerating Medicines Partnership-Alzheimer’s Disease (AMP-AD) MayoRNAseq project [4]. We used the synapse command-line client to download data files using the command:

~~~
SELECT * FROM syn11346063 WHERE ((analysisType=‘differentialExpression’))
~~~

We retained 26 differential gene expression data files from the MayoRNAseq study (13 temporal cortex, 13 cerebellum).

#### Coverage analysis pipeline

We collected intermediate analysis files from a depth-of-coverage analysis pipeline previously applied to several thousand genomes [5]. In particular, we selected for analysis:

- totalCoverageByGC.out: a high-level summary of observed coverage levels stratified by chromosome and by G+C content bins (29 lines per file)
- normalizedCoverage1000.out.gz: the normalized coverage in 1 kb windows (~2.86 million lines)
- HMMSegCoverage1000.out.gz: a segmentation of the genome into ranges of uniform ploidy (~1350 lines)
- filteredCNVs1000.out.gz: a list of genomic ranges with unusual ploidy relative to a reference population (300-400 lines)

## Author contributions

GG conceived of the study. GG, RK, EWD designed and implemented the system. GG, EWD performed analyses. GG, RK, EWD wrote the manuscript. All authors edited and approved its final Version.

## Acknowledgements

This work was supported by NIH grant U54 EB020406.

The Mayo RNAseq study data was led by Dr. Nilüfer Ertekin-Taner, Mayo Clinic, Jacksonville, FL as part of the multi-PI U01 AG046139 (MPIs Golde, Ertekin-Taner, Younkin, Price) using samples from the following sources:

■ The Mayo Clinic Brain Bank. Data collection was supported through funding by NIA grants P50 AG016574, R01 AG032990, U01 AG046139, R01 AG018023, U01 AG006576, U01 AG006786, R01 AG025711, R01 AG017216, R01 AG003949, NINDS grant R01 NS080820, CurePSP Foundation, and support from Mayo Foundation.
■ Sun Health Research Institute Brain and Body Donation Program of Sun City, Arizona. The Brain and Body Donation Program is supported by the National Institute of Neurological Disorders and Stroke (U24 NS072026 National Brain and Tissue Resource for Parkinson’s Disease and Related Disorders), the National Institute on Aging (P30 AG19610 Arizona Alzheimer’s Disease Core Center), the Arizona Department of Health Services (contract 211002, Arizona Alzheimer’s Research Center), the Arizona Biomedical Research Commission (contracts 4001, 0011, 05-901 and 1001 to the Arizona Parkinson's Disease Consortium) and the Michael J. Fox Foundation for Parkinson’s Research.

## References

1. Manimaran S, Selby HM, Okrah K, Ruberman C, Leek JT, Quackenbush J, et al. BatchQC: interactive software for evaluating sample and batch effects in genomic data. Bioinformatics. 2016;32:3836–8.

2. Ward J, Cole C, Febrer M, Barton GJ. AlmostSignificant: simplifying quality control of high-throughput sequencing data. Bioinformatics. 2016;32:3850–1.

3. Ewels P, Magnusson M, Lundin S, Käller M. MultiQC: summarize analysis results for multiple tools and samples in a single report. Bioinformatics. 2016;32:3047–8.

4. Allen M, Carrasquillo MM, Funk C, Heavner BD, Zou F, Younkin CS, et al. Human whole genome genotype and transcriptome data for Alzheimer’s and other neurodegenerative diseases. Sci Data. 2016;3:160089.

5. Glusman G, Severson A, Dhankani V, Robinson M, Farrah T, Mauldin DE, et al. Identification of copy number variants in whole-genome data using reference coverage profiles. Front. Genet. 2015;5:1–13.

6. Modern Visualization for the Data Era [Internet]. [cited 2018 Feb 1]. Available from: https://plot.ly/

